# General stain deconvolution of histopathology images with physics-guided deep learning

**DOI:** 10.1101/2022.12.06.519385

**Authors:** Jianan Chen, Lydia Y. Liu, Wenchao Han, Dan Wang, Alison M. Cheung, Hubert Tsui, Anne L. Martel

## Abstract

Advances have been made in the use of deep learning to extract quantitative and predictive information from digital pathology slides, yet many barriers remain before clinical translation and deployment. In particular, models need to be generalizable despite the wide variations in image characteristics due to inter-scanner variability and differences in slide preparation protocols. This has led to an interest in stain deconvolution methods that could correct for the variability in image appearances. However, most existing stain deconvolution methods were developed and validated on specific datasets and perform poorly on unseen data. We developed Physics-Guided Deep Image Prior network for Stain deconvolution (PGDIPS), a method that combines a novel optical physics model and a self-supervised deep neural network to perform deconvolution for various classes and any number of stains, without the need of training data. PGDIPS outperformed state-of-the-art approaches for the deconvolution of conventional stain combinations, enabled analysis of previously unsupported special stains, and provided superior interpretability by explicitly encoding representations for stain properties and the light transmittance/absorbance process. PGDIPS is publicly available as an end-to-end off-the-shelf tool that does not require data curation, domain knowledge or high computation power.

## Main

Digital pathology is a rapidly growing field where stained tissue samples imaged by scanning devices at microscopic scales are digitized so that they can be analyzed by experts and increasingly, machine learning models^1,2^. Computational pathology aims to extract quantitative features from digital pathology data, often complemented with clinical information and data from other omics domains, in order to improve disease diagnosis, treatment selection and drug target discovery^3–5^. Many hurdles remain to be overcome before advances in computational pathology can be translated into routine use^6^. In particular, models need to be generalizable despite the wide variation in image characteristics due to inter-scanner variability and differences in slide preparation protocols^7^. This has led to an interest in stain deconvolution (SD) methods that aim to correct for the variability in image appearances. Despite being a key pre-processing step in computational pathology, stain deconvolution remains an open problem^8^.

Conventionally, SD is formulated as a matrix decomposition problem for deconvoluting the optical density of stained slides into color vectors that represent the spectral absorptivity of the stained structure for each type of stain, and concentration maps that reflect the spatial distribution of strained structures^9^. Conventional SD methods are based on the Beer-Lambert Law, which relates, linearly, the attenuation of non-scattering light to the properties of the stained tissue through which the light is travelling^10^. Application of Beer’s law is hindered under certain conditions due to complex non-linear image formation, including but not limited to saturation effect from high concentrations, scatter effects, and light polychromaticity, which are ubiquitous conditions in digital pathology images^11^. This leads to systemic deconvolution error, especially for special stains, in conventional SD algorithms^12,13^. Problem-specific SD algorithm design, a time-consuming process that requires extensive domain knowledge, method development expertise and large datasets, is often required for the analysis of non-routine stains^14^.

Neural-network-based SD approaches have been proposed to alleviate the shortcomings of traditional SD methods^15^. Although the increased flexibility of deep-learning-based SD models bypasses the limitations of the Beer-Lambert Law, it is often at the expense of accessibility and generalizability. While a singly-stained slide is considered the gold standard for ground truth, few networks are trained this way because it is impractical to collect singly-stained images for every stain^8^. Thus, most existing deep-learning-based approaches are unsupervised but instead depend on large, carefully curated datasets and demand a high level of expertise in network design and training for each dataset of interest^16–18^. This data-driven nature of existing deep-learning-based SD methods makes them extremely powerful on the datasets they have been trained on, but also stomps their generalizability on unseen data.

State-of-the-art conventional SD algorithms such as Macenko^19^ and Vahadane^20^ are still the go-to methods for general SD tasks, despite the limited generalizability and systematic errors in these approaches^21^. To address the lack of a general SD algorithm in the field, we developed Physics-Guided Deep Image Prior network for Stain deconvolution (PGDIPS), an end-to-end stain deconvolution algorithm that can robustly and accurately deconvolve arbitrary numbers and types of stains. PGDIPS encodes an adapted physics model that includes additional parameters describing background illumination and non-linearities in light transmission, providing superior interpretability. By utilizing deep image prior (DIP), a neural network training scheme that learns the building blocks of any given image solely from reconstructing the image itself^22^, PGDIPS performs stain deconvolution on the fly with no training data, and can thus be applied to special stains where large quantities of data are not readily available. PGDIPS is also uniquely accessible: image processing steps including background illumination correction, color vector estimation and stain deconvolution are performed fully end-to-end and off-the-shelf on entry-level graphics processing units. In this report, we validated PGDIPS on different combinations of digital pathology stains and benchmarked PGDIPS against state-of-the-art SD algorithms to demonstrate its superior applicability and performance.

## Results

### PGDIPS algorithm

PGDIPS implements an end-to-end neural network for the deconvolution of digital pathology images (**Figure 1a**). PGDIPS represents a new paradigm for general stain deconvolution (SD) that combines an optical physics model and a self-supervised deep learning model. The network encodes the optical physics model in its structure to leverage prior knowledge, then follows a deep image prior (DIP) training scheme to solve the optical physics model and perform stain deconvolution without the need for training data.

**Figure 1:**
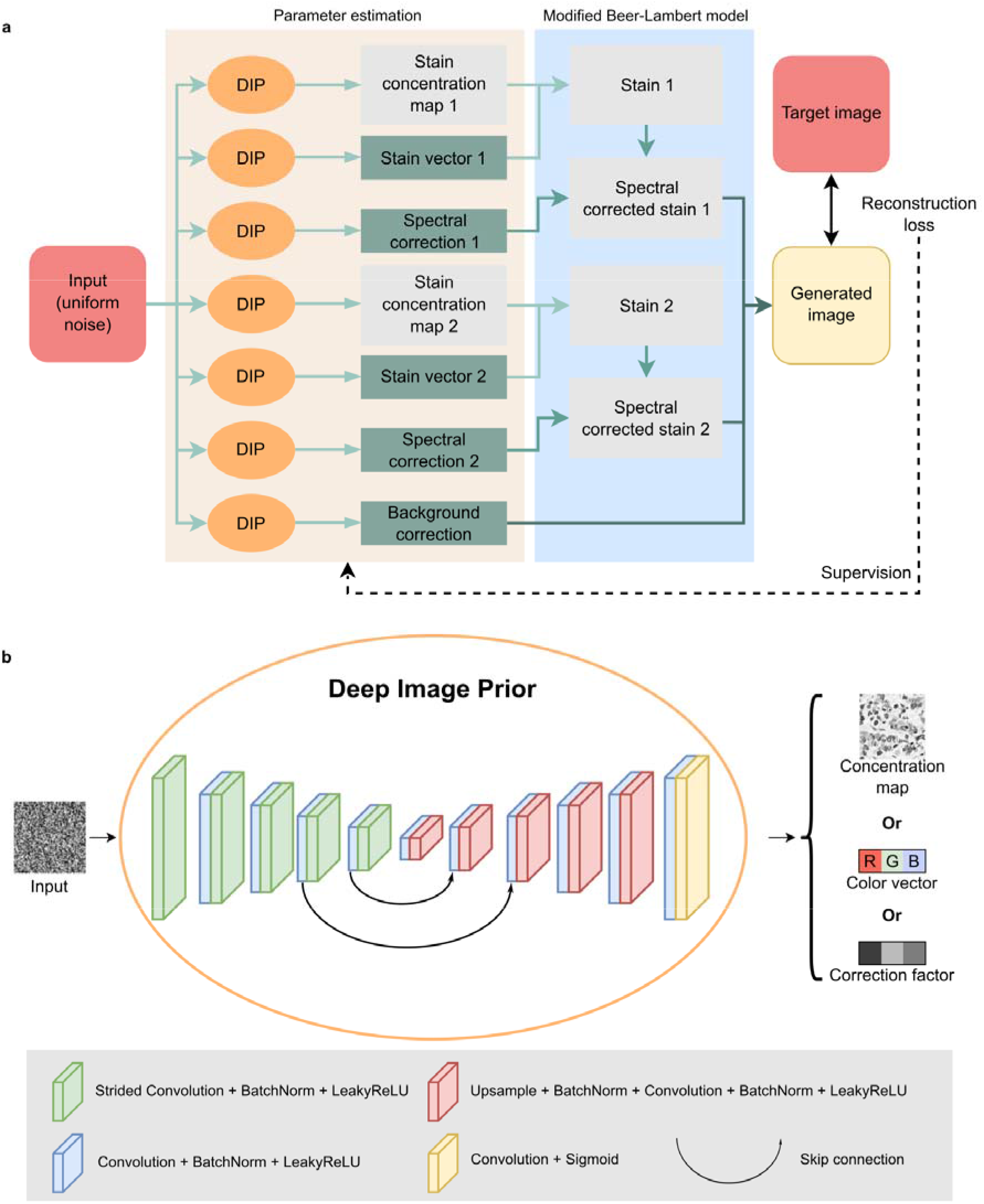
PGDIPS network structure. **a**) PGDIPS network structure diagram featuring a modified Beer-Lambert model solved by components estimated by multiple deep image prior (DIP) modules. A random noise image of the same size as the target image is first generated as network input. The noise image is fed into multiple DIP modules, each responsible for generating a concentration map, stain color vector or correction factor. The generated concentration maps and parameters are combined using the modified Beer-Lambert model to reconstruct the target image. The mean absolute error between the reconstruction and the target image is the primary source of end-to-end supervision for the entire network. **b**) Overview of deep image prior network structure. Deep image prior refers to the process of extracting image characteristics from learning to reconstruct an image through a neural network that has limited capacity. Deep image prior networks are implemented as U-shaped encoder-decoder networks (UNets) with skip connections, that take uniform noise as input and generate images or vectors as outputs. Strided convolution with stride of 2 is used to downsample feature maps and upsampling is performed using bilinear interpolation with a factor of 2. Each convolution and strided convolution layer are followed by batch normalization and LeakyReLU activation, except for the last convolution layer in the network, which is followed by a Sigmoid activation.

We first adapted the Beer-Lambert Law for SD by introducing two additional components (**Methods**). The first is a spectral correction factor that aims to adjust, after non-linearity correction, the contribution of each stain on perceived slide appearance (**Extended Data Figure 1**). The second is a background illumination correction factor, in order to account for the characteristic radiation spectrum of the illumination source. We reasoned that these components could alleviate the systemic error caused by discrepancy between the non-linear signal formation and the linear assumption of existing SD approaches and enable background illumination correction to be incorporated seamlessly into SD.

Estimating additional parameters is typically beyond the capability of non-negative matrix factorization and singular value decomposition algorithms, which are the backbone of conventional SD approaches. Therefore, we encoded PGDIPS to solve the adapted optical physics model by utilizing multiple neural networks, each responsible for estimating either a color vector (a vector), a correction factor (a vector) or a stain concentration map (an image) (**Figure 1b, Methods**). The parameters and concentration maps are then combined using the optical physics model to generate a reconstructed version of the target image (*i.e*. the image undergoing stain deconvolution). The mean absolute error between the reconstruction and the target image is then used to guide the training of the entire PGDIPS network. We also included an exclusion loss calculated between stain concentration maps that encourages the unmixing of stains and a color fixing loss calculated between stain color vectors that adjusts the focus of the network from learning colors to stain concentrations in different stages of learning.

In order to achieve true stain generalizability, we constructed PGDIPS to utilize the self-supervised training enabled by DIP modules^22^. The classic DIP network attempts to find a mapping that transforms random noise into the target image while capturing image-intrinsic priors as building blocks for reconstruction (*i.e*. deep image priors) in the convolutional filters of the DIP network. Multiple DIPs have been combined, typically using a simple linear equation, for image decomposition tasks such as segmentation and dehazing^23^. We extended this concept and assembled multiple DIP modules in the manner of our adapted optimal physics model, with the number of DIP modules determined by the number of stains to deconvolute. This network construction has the added advantage of learning solely from the target image, thus circumventing the risk of overfitting.

### Benchmarking and validation of PGDIPS

To evaluate the utility of our proposed paradigm, we applied PGDIPS on digital pathology images with six distinct types of stains (**Figure 2**). In addition to the widely-used Hematoxylin & Eosin (H&E) staining (**Figure 2a**), PGDIPS also deconvoluted the special stains tested with visual correctness, namely hematoxylin & 3,3’-diaminobenzidine (DAB) staining (**Figure 2b**), Alcian Blue staining (**Figure 2c**), anti-fibronectin antibody staining (**Figure 2d**), Masson’s Trichrome staining (**Figure 2e**), and clinical-grade dual IHC anti-SMA and anti-CD34 antibody staining (**Figure 2f**). Since no ground truth was available, we qualitatively compared the outputs of PGDIPS with two state-of-the-art approaches for general SD tasks: Macenko, a SD algorithm that estimates stain color vectors based on singular value decomposition, and Vahadane, a SD algorithm that preserves structure with sparsity-constrained non-negative matrix factorization^19,20^. As with Macenko and Vahadane, PGDIPS handles general SD tasks for images of all dimensions and resolution, and requires no hyperparameter tuning for application to any stains. Unlike Macenko and Vahadane, which both had difficulties with one or more special stains, PGDIPS achieved visually superior reconstructed images with higher fidelity and colour vectors with higher visual accuracy (**Extended Data Figure 2**).

**Figure 2:**
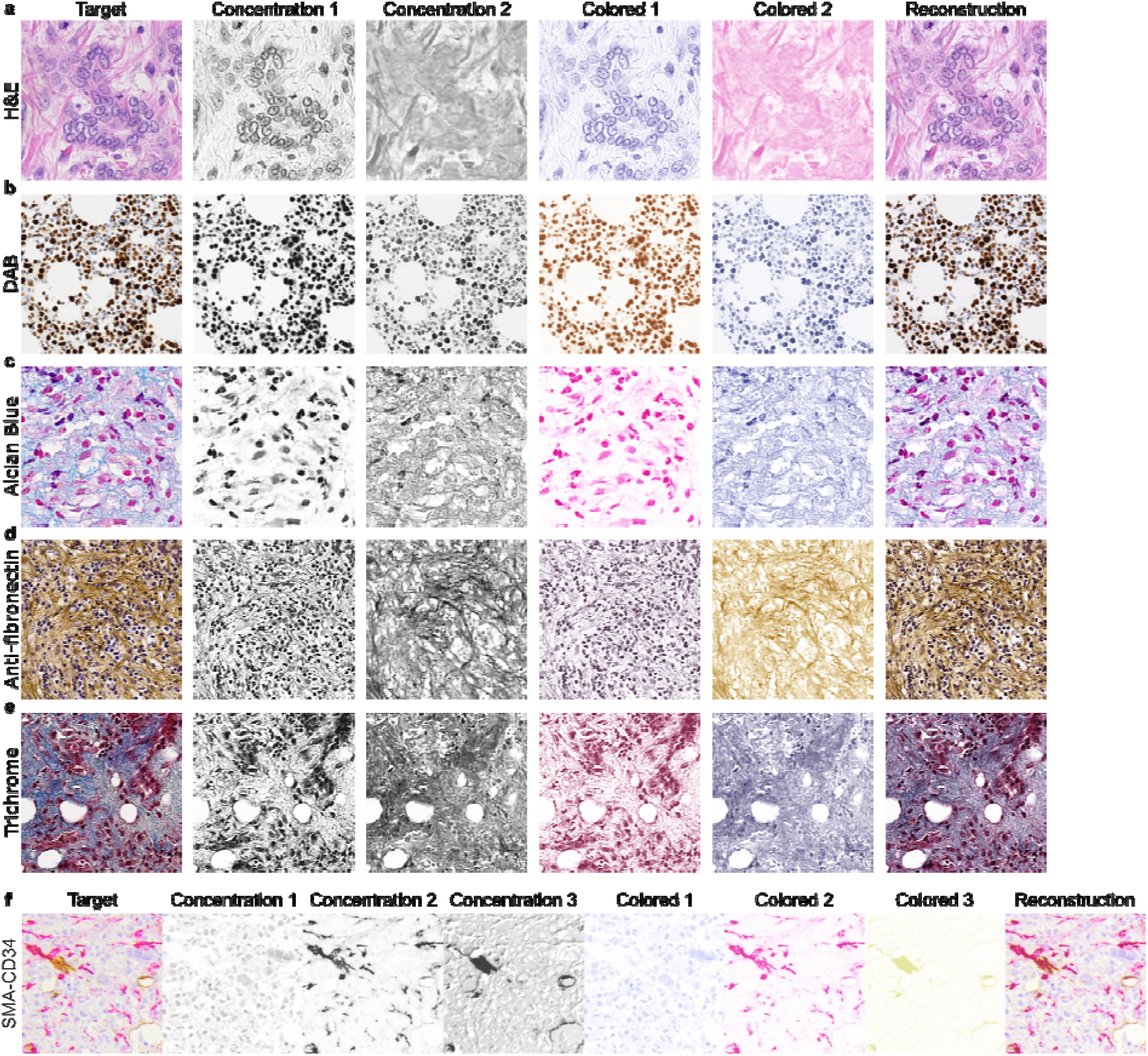
PGDIPS generalizes to different types of stainings. PGDIPS stain deconvolution results of slides stained with **a**) Hematoxylin and Eosin (H&E) **b**) hematoxylin and 3,3’-diaminobenzidine (DAB) **c**) Alcian Blue **d**) anti-fibronectin antibody **e**) Masson’s Trichrome **f**) SMA-CD34 dual antibodies. Columns are organized to display the target image, the stain concentration maps estimated by PGDIPS, the colored concentration maps (matrix multiplication of the corresponding stain color vectors and stain concentration maps) and the reconstructed image.

We next sought to validate and benchmark the performance of PGDIPS quantitatively, first for the most common pathology staining, H&E staining, on an in-house breast cancer pathology dataset with 2008 co-registered H&E and immunofluorescence (mIF) 4’,6-diamidino-2-phenylindole (DAPI) images (**Figure 3a**). Instead of using singly-stained images as the ground truth, we used co-registered mIF images of the same tissue as a surrogate, because singly-stained images are not routinely collected, and the appearance of hematoxylin-only images are also affected by scanner model and background illuminations. mIF images, on the other hand, are increasingly collected in multi-modality studies, and are dark-field images that are immune to the effects of background illuminations. We therefore used DAPI staining as a surrogate label for ground-truth hematoxylin concentration as DAPI and hematoxylin both stain cell nucleus (**Figure 3b**). PGDIPS achieved significantly higher deconvolution accuracy compared to Macenko and Vahadane, measured by Pearson correlation (PRC) between the deconvoluted hematoxylin stain and co-registered DAPI stain (**Figure 3c**; PGDIPS: 0.592 ± 0.116, mean ± standard deviation [sd]; Macenko: 0.553 ± 0.166, *p* < 0.001, *t* = 8.755; Vahadane: 0.468 ± 0.130, *p* < 0.001, *t* = 32.067; *t*-tests). PGDIPS achieved an even higher performance gap in the structural similarity index (SSIM), a measure for tissue structure preservation^24^, suggesting that the image-prior-based design of PGDIPS has higher fidelity in preserving nuclei structures, which may have large implications for downstream analyses (**Figure 3d**; PGDIPS: 0.521 ± 0.123, mean ± sd; Macenko: 0.350 ± 0.150, *p* < 0.001, *t* = 39.434; Vahadane: 0.235 ± 0.132, *p* < 0.001, *t* = 71.171; *t*-tests). To demonstrate the stability of PGDIPS across the cohort of 2008 images, we visualized the estimated hematoxylin and eosin color vectors in the hue saturation value (HSV) color space (**Figure 3e-g**). The compact distribution of color vectors estimated by PGDIPS, as compared to those estimated by Macenko and Vahadane color vectors, reflect the performance stability of PGDIPS in a cohort processed with the same staining protocol and same scanner.

**Figure 3:**
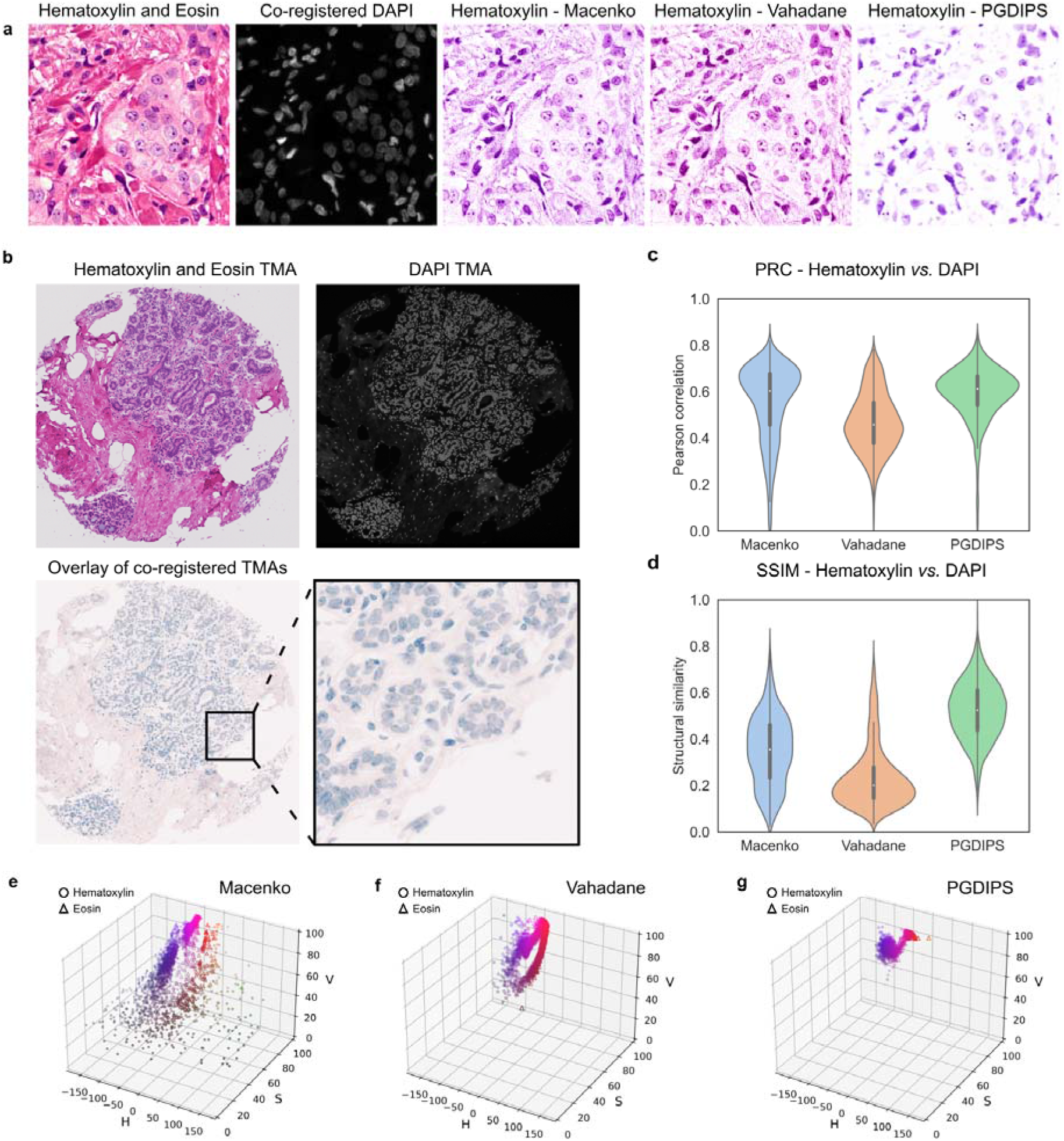
PGDIPS outperforms state-of-the-art algorithms on a co-registered H&E and mIF dataset. **a**) Representative pair of co-registered Hematoxylin and Eosin (H&E) and multiplex immunofluorescence (mIF) 4’,6-diamidino-2-phenylindole (DAPI) images, and the deconvolution results for hematoxylin from Macenko, Vahadane and PGDIPS. DAPI and hematoxylin both stain the nuclei. **b**) Visualization of the co-registration process on a representative tissue microarray (TMA). Images used for deconvolution were cropped patches from co-registered TMAs. Top left: H&E stained breast TMA core; Top right: corresponding mIF DAPI TMA core; Bottom left: DAPI (blue) registered to H&E (light pink), with a zoomed-in patch displayed. **c**) Pearson correlation (PRC) between mIF DAPI and deconvoluted hematoxylin comparison between algorithms. **d**) Structural similarity (SSIM) between co-registered DAPI and deconvoluted hematoxylin comparison between algorithms. Stain color vectors estimated in hue saturation value (HSV) color space by **e**) Macenko **f**) Vahadane **g**) PGDIPS. n = 2008 for all panels.

For validating and benchmarking the performance of PGDIPS on a special stain, we used the test set of DeepLIIF^8^, a co-registered hematoxylin and DAB stain, hematoxylin-only stain and mIF dataset with 598 images (**Figure 4a**). The reconstructed images generated by PGDIPS achieved a closer visual approximation of the target image compared to Macenko and Vahadane (**Figure 4b**), in part due to its capability of separating out the background illumination (**Figure 4c**). PGDIPS also achieved superior performance in deconvolution accuracy and structure preservation for nuclei staining in this special stain, both when using DAPI as the surrogate for hematoxylin concentrations (**Figure 4d-e**; PGDIPS: PRC = 0.768 ± 0.109, SSIM = 0.588 ± 0.117, mean ± sd; Macenko: PRC = 0.751 ± 0.105, *p* = 0.008, *t* = 2.677, SSIM = 0.387 ± 0.065, *p* < 0.001, *t* = 36.615; Vahadane: PRC = 0.693 ± 0.110, *p* < 0.001, *t* = 11.883, SSIM = 0.312 ± 0.056, *p* < 0.001, *t* = 51.955; *t*-tests) and against the hematoxylin-only stain as the gold standard ground truth (**Figure 4f-g**; PGDIPS: PRC = 0.787 ± 0.096, SSIM = 0.456 ± 0.105, mean ± sd; Macenko: PRC = 0.768 ± 0.094, *p* = 0.001, *t* = 3.412, SSIM = 0.302 ± 0.078, *p* < 0.001, *t* = 28.671; Vahadane: PRC = 0.707 ± 0.103, *p* < 0.001, *t* = 13.795, SSIM = 0.244 ± 0.068, *p* < 0.001, *t* = 41.411; *t*-tests). We similarly treated mIF Ki67 as the label for evaluating the deconvolution of DAB stains, as they both stain Ki67-positive cells and DAB-only stains were not available. The performance of PGDIPS on deconvolution accuracy and cell structure preservation was maintained for the special DAB stain, and significantly surpassed that of Macenko and Vahadane (**Figure 4h-i**; PGDIPS: PRC = 0.716 ± 0.133, SSIM = 0.767 ± 0.095, mean ± sd; Macenko: PRC = 0.666 ± 0.130, *p* < 0.001, *t* = 6.550, SSIM = 0.401 ± 0.039, *p* < 0.001, *t* = 87.012; Vahadane: PRC = 0.672 ± 0.136, *p* < 0.001, *t* = 5.693, SSIM = 0.334 ± 0.032, *p* < 0.001, *t* = 105.181; *t*-tests). Color vectors estimated by Macenko showed a consistent bias that led to spurious concentrations in the deconvoluted DAB layer and while Vahadane achieved a more accurate color deconvolution, it did not preserve cell structures in the deconvoluted hematoxylin layer (**Figure 4b**). This highlighted the challenges in special stain deconvolution and validated the generalizability of PGDIPS, especially given the larger stain variability in DeepLIIF test set due to the differential presence of Ki67-positive cells in images, which resulted in the larger variation in color vectors in the HSV space, as well as the lower long tail in PRC scores across all algorithms (**Figure 4j-l, Extended Data Figure 3**).

**Figure 4:**
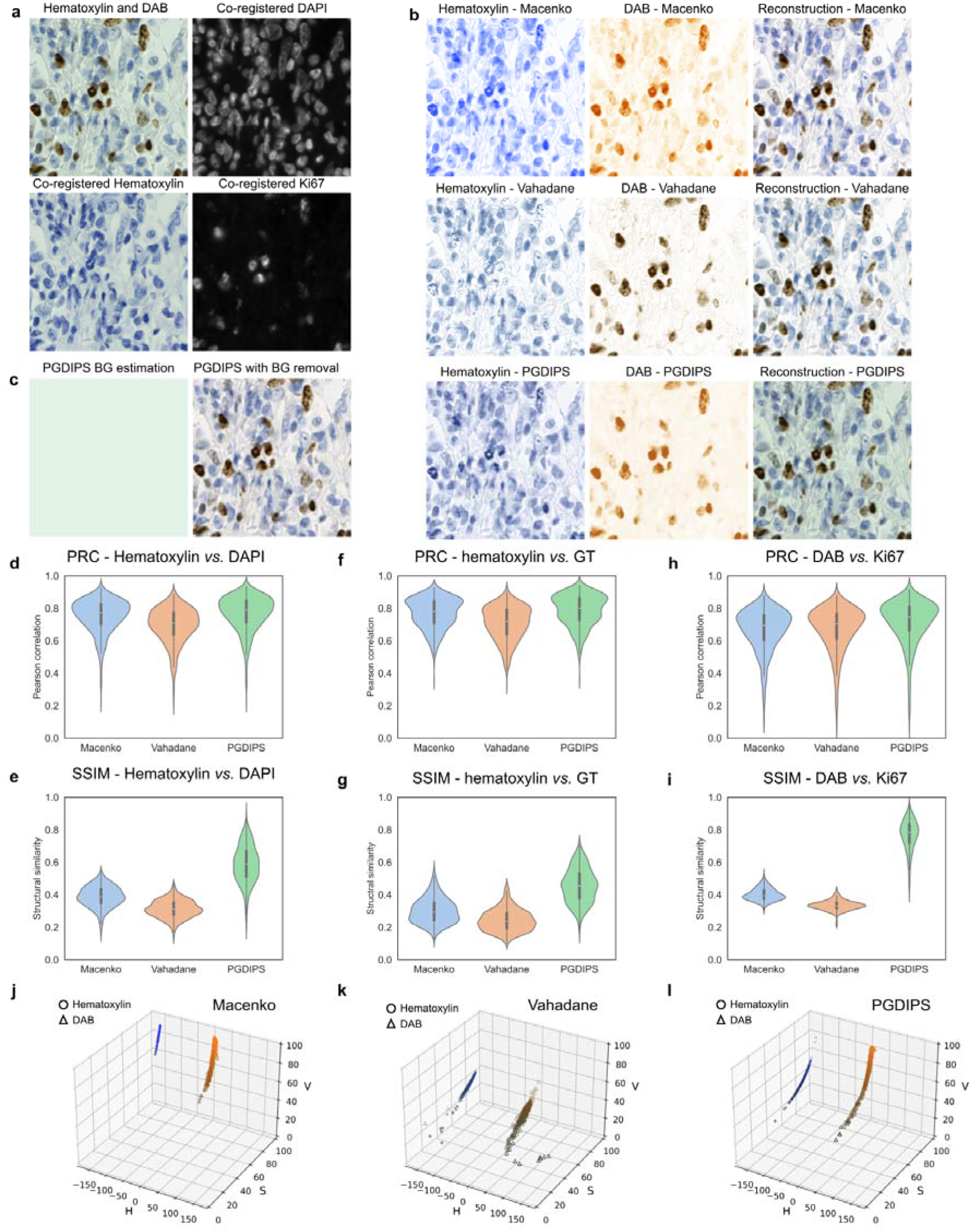
PGDIPS outperforms state-of-the-art algorithms on a co-registered IHC and mIF dataset. **a**) Representative tissue with co-registration of immunohistochemistry (IHC) 3,3’-diaminobenzidine (DAB) with hematoxylin as counterstain and multiplex immunofluorescence (mIF) of 4’,6-diamidino-2-phenylindole (DAPI) and Ki67, along with a corresponding co-registered hematoxylin-only-stained image. **b**) Hematoxylin and DAB staining deconvolution results from Macenko, Vahadane and PGDIPS. **c**) Background (BG) illumination estimated by PGDIPS and a background corrected version of the reconstructed image generated by PGDIPS. mIF DAPI and deconvoluted hematoxylin **d**) Pearson correlation (PRC) **e**) Structural similarity (SSIM) comparison between algorithms. Hematoxylin-only ground truth (GT) and deconvoluted hematoxylin **f**) PRC **g**) SSIM. mIF Ki67 and deconvoluted DAB **h**) PRC **i**) SSIM. Stain color vectors in hue saturation value (HSV) color space estimated by **j**) Macenko **k**) Vahadane **l**) PGDIPS. n = 598 for all panels.

### Down-stream application

As a direct down-stream application of SD, digital pathology images can be normalized to apply an uniform style to images acquired through different scanners and removes a source of variability from down-stream analyses, such as cell segmentation or image classification tasks^25,26^. This is typically achieved by applying a standard set of stain color vectors from the target stain type to all estimated concentration maps^20,27^. We extended this approach by not only applying the deconvoluted color vectors of the reference image to the deconvoluted concentration map of the source image, but also applying the PGDIPS-estimated spectral correction factors and background illumination factor from the reference image, with the reasoning that this should achieve higher normalization fidelity. Visually, stain normalization using PGDIPS achieved a consistent appearance in the MIDOG2022 dataset with a randomly selected reference image, across different studies, scanners and tissue types, again demonstrating the accuracy of parameter inference by PGDIPS (**Figure 5**). These additional parameters can also be plugged into other SD methods as additional constraints or components for matrix decomposition for improving their performance in SD and downstream tasks.

**Figure 5:**
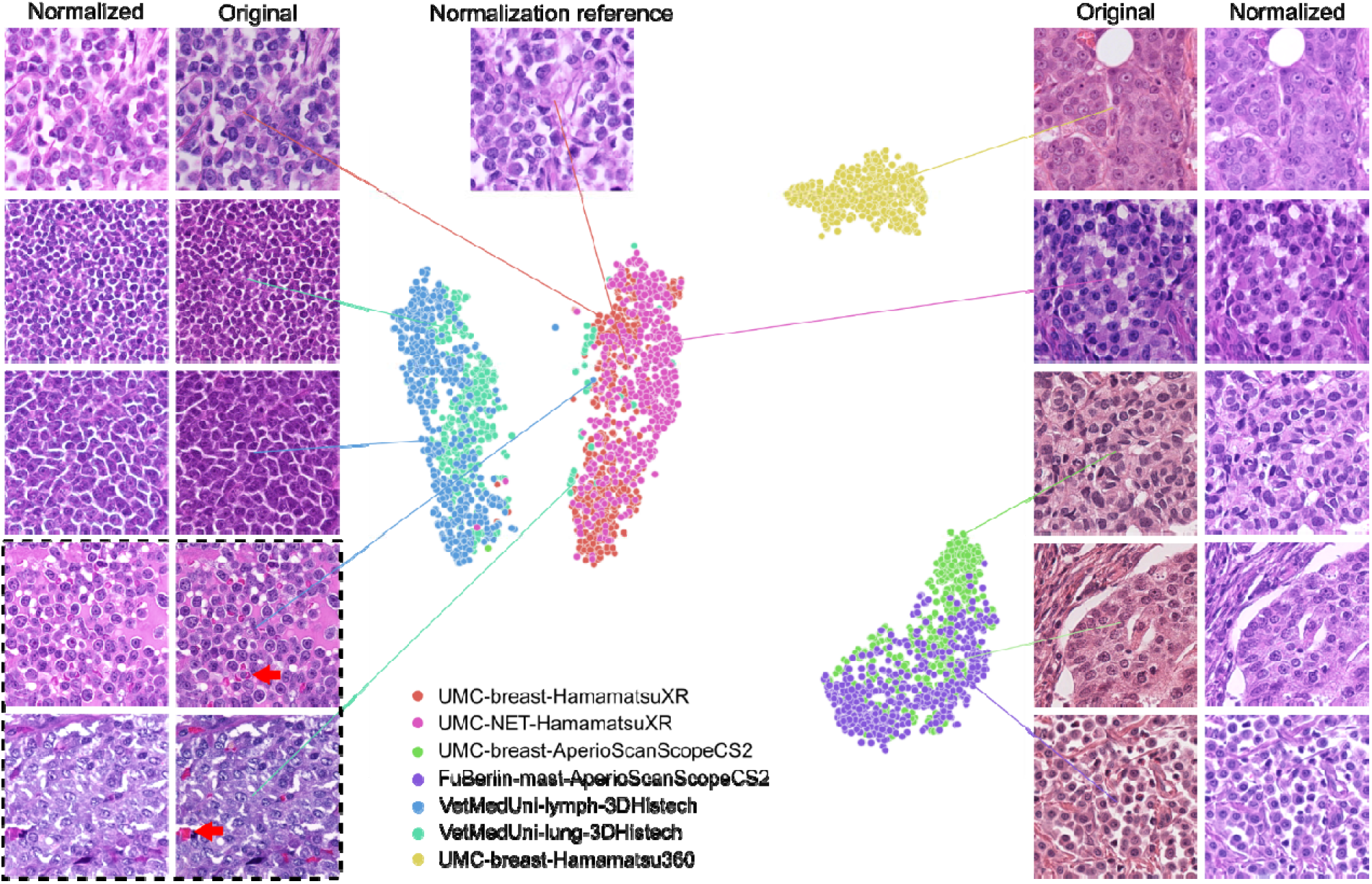
PGDIPS-estimated physical parameters cluster by scanner type. Two-dimensional representations of estimated physical parameters of Hematoxylin and Eosin (H&E) images taken from 7 different studies in the MIDOG2022 dataset, estimated by t-distributed stochastic neighbor embedding (t-SNE, n = 3186). Representative images from each cluster are shown with their normalized versions generated by PGDIPS, where the target normalization parameters were generated by applying PGDIPS on a reference image randomly selected from the dataset. Most samples were correctly clustered by scanner type, and representative cluster misassignments are shown in the bottom left (black dotted box). The artifacts (red arrows) were likely the cause of misassignment. Breast: human breast cancer; NET: human neuroendocrine tumor; Lymph: canine lymphoma; Lung: canine lung cancer; Mast: canine cutaneous mast cell tumor. UMC: University Medical Center Utrecht; VetMedUni: University of Veterinary Medicine Vienna; FuBerlin: Free University of Berlin.

## Discussion

We present PGDIPS, an algorithm for zero-shot general stain deconvolution. PGDIPS outperformed existing approaches for SD in a variety of stain types and is, to our knowledge, the first SD method that is capable of deconvolving any type of stain without task-specific adaptation. The strong generalizability of PGDIPS arises from the synergy between the encoded optical physics model and the architecture of the deep neural network. Specifically, the neural network leverages the regulation of the physics model to achieve optimization solely based on the target image itself, which is a novel approach to SD. This not only avoids the labor-intensive and time-consuming data collection and curation process, but also prevents overfitting to the training distribution. The DIP modules in PGDIPS further ensures that the image structures of the target image are preserved in the deconvoluted stain layers, as the limited number of convolution filters in DIP modules are forced to capture the intrinsic image-statistics of the target image. This can have a significant impact on downstream tasks such as cell segmentation, disease classification and outcome prediction, where accurate feature extraction is key.

PGDIPS also introduces a higher level of interpretability to stain deconvolution. We have configured PGDIPS to output the list of parameters estimated by each DIP module, which describes conditions and properties involved in the process of light transmittance/absorbance, in contrast to most deep-learning-based SD algorithms that only output normalized images. By introducing a stain spectral correction factor that can only be estimated with deep learning, we were able to broaden the application scope of the Beer-Lambert Law to stains where the simplifying assumptions made by existing deconvolution approaches do not hold, as evident in the significantly reduced deconvolution errors and more accurate stain color estimations. PGDIPS also models the properties of the background light which allows background illumination correction to be integrated into the color deconvolution process, as opposed to as a separate pre-processing step^28^.

Stain deconvolution with PGDIPS fully relies on information supplied by the target image. As a result, the performance of PGDIPS may deteriorate when the target image quality is low, especially when staining is not consistent throughout the target image, or when the absorption spectra of the stains or the background illumination are too similar. For example, when there is a high-concentration ink-mark in the image, PGDIPS will tend to compute a biased color vector and concentration map as a compromise to fit both the stained structures and the ink-mark. Similarly, PGDIPS will have suboptimal performance in assigning structures between similar stains. These remain universal challenges for stain deconvolution and could be most easily addressed with better quality control in slide preparation protocols. PGDIPS offers potential solutions to these challenges: by stacking more DIP modules and updating the physics model to take large artifacts into account, and by estimating and interpreting physics parameters to inform slide preparation quality control in real time.

Stain deconvolution also serves as a prerequisite step for stain quantification tasks because deconvoluted stain layers simplify cell segmentation and counting^29^. We also developed a stain analysis tool, which takes parameters estimated by PGDIPS and automatically performs SD and stain quantification for whole slide images. We have released the tool as a plugin for Sedeen Viewer, a freely available pathology viewer^30^.

In summary, we anticipate that PGDIPS will usher in a new generation of stain deconvolution methods with strong generalizability across a wide variety of stains without the need for expert knowledge or data collection. We have made PGDIPS an easily accessible off-the-shelf tool for state-of-the-art general-purpose stain deconvolution that could be incorporated into any digital pathology workflow.

## Methods

### Image Data

Zero-shot stain deconvolution was performed on a Hematoxylin and Eosin (H&E) stained breast tissue image from the MIDOG dataset, a 3,3’-diaminobenzidine (DAB) stained bone marrow image from our in-house immunohistochemistry (IHC) acute myeloid leukemia cohort, and an Alcian blue image, a Masson’s Trichrome image and an anti-fibronectin antibody IHC image from our in-house Hodgkin’s lymphoma cohort (Bahlmann, L. C. *et al*., under review), and a SMA-CD34 dual IHC (alkaline phosphatase for SMA and DAB for CD34 with hematoxylin counterstain) bone marrow image from our in-house myeloproliferative neoplasms cohort^31^. All the images were randomly cropped as 512×512 patches from a whole slide image.

The in-house dataset of breast tissue microarrays (TMAs) with H&E and 4’,6-diamidino-2-phenylindole (DAPI) mIF staining of the same tissue was scanned using a Zeiss Axiovision Z1 motorized stage microscope (GE Research Niskayuna NY), as previously described^32,33^. 2232 non-overlapping 512×512 H&E patches were cropped from 81 TMAs and patches with lower than 10th quantile entropy were discarded to exclude patches with too much background, resulting in 2008 patches. Co-registered DAB and mIF images were from the test set of DeepLIIF^8^, where 598 sets of images scanned using a ZEISS Axioscan scanner were used as is. The training set of MIDOG 2022 was also used^34^, including 354 high-power field images generated by imaging 150 human breast cancer cases (3 scanners with 50 cases each: Hamamatsu XR nanozoomer 2.0, Hamamatsu S360 [0.5 NA], Scanner 3: Aperio ScanScope CS2), 55 human neuroendocrine tumor cases (Hamamatsu NanoZoomer XR), 44 canine lung cancer cases (3DHistech Pannoramic Scan II), 55 canine lymphoma cases (3DHistech Pannoramic Scan II) and 50 canine cutaneous mast cell tumor cases (Aperio ScanScope CS2). We randomly selected nine connected non-background 512×512 patches from each high-power field image to generate 3186 patches for the experiment.

DAPI mIF images were co-registered with fixed H&E images in our in-house dataset using scale-invariant feature transform^35^ followed by affine transformation. For the DeepLIIF dataset, mIF images and the hematoxylin and DAB images were also co-registered using affine transformation, as previously described^8^.

### Optical physics model

Following conventional SD methods^9,19^, PGDIPS converts RGB intensities of the target image intensities ***I*** into corresponding optical density values ***OD*** by taking a logarithm operation based on ***I*** normalized by maximum Intensity ***I**_0_* (**Equation 1**). The Beer-Lambert Law then decomposes the optical density values ***OD*** into individual concentration maps ***C*** and stain color vectors ***S*** for each stain (**Equation 2**).

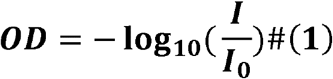

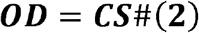

PGDIPS proposes an adapted Beer-Lambert Law as shown in **Equation 3**, where *N* is the number of stains.

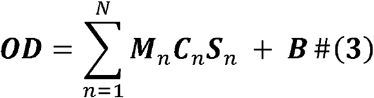

We built upon Haub and Meckel’s work^13^ and addressed the problem of signal non-linearity by proposing a stain spectral correction factor ***M*** to summarize the non-linearities in signal formation and approximating this coefficient using deep image prior modules. For each particular stain, spectral variations such as spectral transmittance, imaging device spectral sensitivity and light source spectral intensity distribution by definition all vary at different wavelengths, making the precise computation of ***OD*** intractable. However, we reason that the main source of deconvolution error comes from the difference in spectral variations between different stains. For example, DAB is known to be a scatterer of light compared to hematoxylin^36^, but in conventional SD models DAB and hematoxylin are estimated using the same linear model. Adding a learnable weight ***M*** allows the deconvolution model to adjust the contribution of each stain and reduce errors from non-linearities.

In addition, PGDIPS estimates the effect of background illumination by denoting the change in optical density values in each channel with a background illumination correction factor ***B***. Background illumination may affect the perceived color of images captured by a scanner, and the effect varies between different scanners. Existing methods rely on an extra pre-processing step based on Gaussian denoising filters or sampling from background pixels to restore the pristine image^37^. PGDIPS integrates the estimation of background illuminations into the generation/reconstruction of the target image, which not only produces an end-to-end SD process, but also minimizes the denoising error as a part of the objectives of the network.

In all reported experiments, ***M*** is defined as a float number for each stain, and ***B*** is defined as a 3-element vector for each image. Both parameters were assumed to be constant across a target image to simplify computation. Estimating ***M*** and ***B*** as images (*i.e*. 1-element vectors and 3-element vectors that vary at each image pixel) may further reduce deconvolution error, but we have found the current deconvolution performance with simplified ***M*** and ***B*** estimation satisfactory.

### Network structure and Deep Image Prior modules

The structure of PGDIPS features multiple Deep Image Prior (DIP) modules assembled under the regulation of the proposed optical physics model. Each DIP module is responsible for one variable regardless of the size of the variable, including stain color vectors (1, 3; dimension 1, dimension 2), stain concentration maps (input dimension 1, input dimension 2), stain spectral correction factors (1, 1) and background illumination (1, 3). For DIP modules that were designed to estimate a vector, the modules were encoded to reconstruct the central n-length vector of the target image instead of the whole image (n = 3 for stain color vector and background illumination, n = 1 for spectral correction factor). In a standard two-color stain deconvolution problem, PGDIPS used 2 DIPs for learning concentration maps (1 per stain), 2 DIPs for estimating stain vectors, 2 DIPs for learning the spectral correction factors, and 1 DIP for estimating background illumination. For images stained with more than two dyes, the network could be easily extended by adding three additional DIP modules per stain (1 concentration map, 1 stain vector and 1 stain spectral correction factor for each additional stain).

Each DIP module followed the design of U-Net, a U-shaped architecture consisting of a downsampling encoder and an upsampling decoder^38^. The input image was uniform noise in the range of 0.5 to 0.5 and the convolution filter size was 5×5. Augmentations consisting of 90-degree rotations and flipping on the input random noise were also used to help the network learn a pose-invariant deep image prior. The encoder consisted of 5 strided convolution blocks for downsampling, where the first block was a convolution layer with a stride of (2,2) and the other blocks consisted of a convolution layer with a stride of (1,1) followed by a convolution layer with a stride of (2,2). The decoder consisted of 5 upsample blocks each composed of a convolution layer with a stride of (1,1), a bilinear upsampling layer with batch normalization and another convolution layer with a stride of (1,1). The scale factor for upsampling was 2 and the input of each layer was padded accordingly. The 4th and 5th strided convolution blocks of the encoder were also skip-connected to the output of the 2nd and 1st upsampling blocks of the decoder, respectively. All convolution layers in the encoder and decoder were followed by batch normalization and LeakyReLU activation. One additional convolution layer followed the encoder and decoder, and was followed by Sigmoid activation.

### Objectives

Given a target image, the main objective of PGDIPS was to train a model ***F***(.) parameterized by a neural network Θ that generates an image based on the optical physics model and minimizes the pixel-wise difference between the generated image and the target image (**Equation 4**).

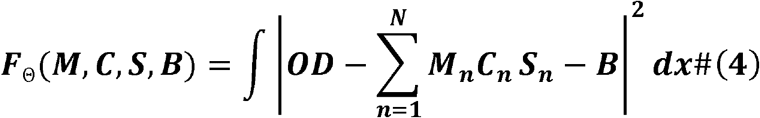

Θ = {*θ*_1_, *θ*_2_,…, *θ*_*K*_} denotes the weights of a series of DIP modules and *x* denotes each pixel. The loss function of PGDIPS was built upon this main objective, with two additional components (**Equation 5**):

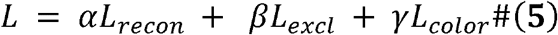

1. Reconstruction loss (recon): L1-losses for optical density (OD) in each channel of the color space (red, green, blue; RGB) and sum of OD across the three channels, between the reconstructed image and the target image, for minimizing generation error.
2. Exclusion loss (excl): exclusion losses between the estimated concentration maps, to encourage separation between different stain layers since different stains should be highlighting different regions of interest.
3. Color fixing loss (color): L2-losses between the estimated stain color vectors and initial stain color vectors (a hyperparameter), for stabilizing color vectors in the early stages of training so that the network focuses on learning internal structure patterns before learning colors.

The weights for each loss were set to ***α*** = 1, ***β*** = 0.01, ***γ*** = 1 to balance the three objectives. The implementation of exclusion loss followed that of prior applications^23,39^. The color fixing loss regulated the network in the first 25% of epochs and was removed subsequently. For standard two-color stain separation, the final loss function consisted of four reconstruction losses, one exclusion loss and two color-fixing losses. The loss functions could be extended for ***N*** ≥ 3 colors by incorporating four reconstruction losses, *N*(*N* – 1)/2 exclusion losses and *N* color-fixing losses.

### Implementation details

The Adam optimizer with *β*_1_ = 0.9, *β*_2_ = 0.999 and an initial learning rate of 0.0005 was used for PGDIPS. A single optimizer was used for updating the parameters of all DIP modules in the network simultaneously. Network parameters were initialized using the default initialization method in PyTorch^40^, namely Kaiming Uniform initialization. Optimization for an image with a size of (512, 512) pixels converged in approximately 4000 steps, and took around 13 minutes for images with 2 stains and 20 minutes for images with 3 stains on a Nvidia RTX 3080 GPU.

PGDIPS was implemented in Python 3.7.9 using PyTorch framework 1.8.1. Python libraries that were also used include torchvision 0.9.1, scikit-image 0.17.2, scikit-video 1.1.11, SciPy 1.6.0, Pillow 8.1.0, NumPy 1.20.0, matplotlib 3.3.2 and pandas 1.2.1. For a complete list of dependencies, please refer to the online documentation available on the GitHub page, available upon publication.

Macenko, Vahadane, PGDIPS were run with default parameters, except for adapting the number of stains. Macenko and Vahadane were implemented using scripts from https://github.com/wanghao14/Stain_Normalization. Default initial color vectors of [0.60, 0.75, 0.29] for the first stain and [0,21, 0.91, 0.36] for the second stain were chosen for calculating the color fixing loss in all experiments with PGDIPS.

### Stain normalization and transformation

Applying PGDIPS for stain normalization followed the standard workflow in the field. Deconvolution was performed for the reference image and all images that needed to be normalized. Then the stain color vectors of all images were replaced by the stain color vector of the reference image to align image appearance while maintaining image content. Stain normalization with PGDIPS involved the additional step of replacing the spectral correction factors and background correction factor of each image with those from the reference image.

### Stain deconvolution on whole slide images

PGDIPS supported stain deconvolution for an image with size up to (1352, 1352) pixels on a NVIDIA RTX 3080 GPU with 10GB of VRAM. In order to apply PGDIPS to a whole slide image (WSI), patches must be extracted, deconvoluted and stitched back together. Alternatives for this process were developed in the interest of runtime (**Supplementary Table 1**). One of the alternatives, which takes around 15 minutes for a WSI, is supported by a stain analysis plugin developed for Sedeen Viewer^30^ (v5.5.0 onwards).

### Data visualization

Box plot elements were reported with the miniature box plots inside violin plots (black box: quartile one [Q1] to quartile three [Q3], white dot: median, whiskers: lowest or highest values within 1.5x the interquartile range from Q1 or Q3, respectively). The hue saturation value (HSV) color space was chosen over the RGB color space for visualizing color vectors because intra-dataset color variations were expected to be present mostly in the saturation and value (brightness) channels. Visualization of estimated physical parameters was performed with t-distributed stochastic neighbor embedding (t-SNE), using n-component of 2, perplexity of 75 and 5000 iterations. All statistical tests were two-sided unless otherwise stated. All t-tests assumed equal variances of the two groups unless otherwise stated.

## Data Availability

The co-registered breast cancer TMA H&E and mIF dataset is available from the corresponding author upon reasonable request. DeepLIIF is publicly available at https://zenodo.org/record/4751737#.YV379XVKhH4. MIDOG is publicly available at https://imig.science/midog/the-dataset/. Source data, including the images used for generating Figure 2, are provided with this paper.

## Code Availability

All code was implemented in Python using Pytorch as the primary deep learning package. All code and scripts to reproduce the experiments of this paper are available at https://github.com/GJiananChen/PGDIPS.

## Author Contributions

Study Design: JC, ALM

Pipeline and Method Development: JC

Cohort Development, Pathology Review, Sample Processing: WH, DW, AMC, HT

Data Analysis: JC, LYL

Supervised Research: ALM

Prepared the manuscript with input from all co-authors: JC, LYL, ALM

Reviewed and Approved Text: all authors

## Acknowledgements

The authors thank the software support team at Sunnybrook Research Institute for their support in developing and deploying the stain analysis tool. We would also like to thank Dr. Martin Yaffe, Dr. Molly S. Shoichet, and Laura C. Bahlmann for providing data. We thank Dr. Jun Ma for constructive comments, and Dr. Steve Leonard for his insightful questions.

## Extended Data Figures

**Extended Data Figure 1:**
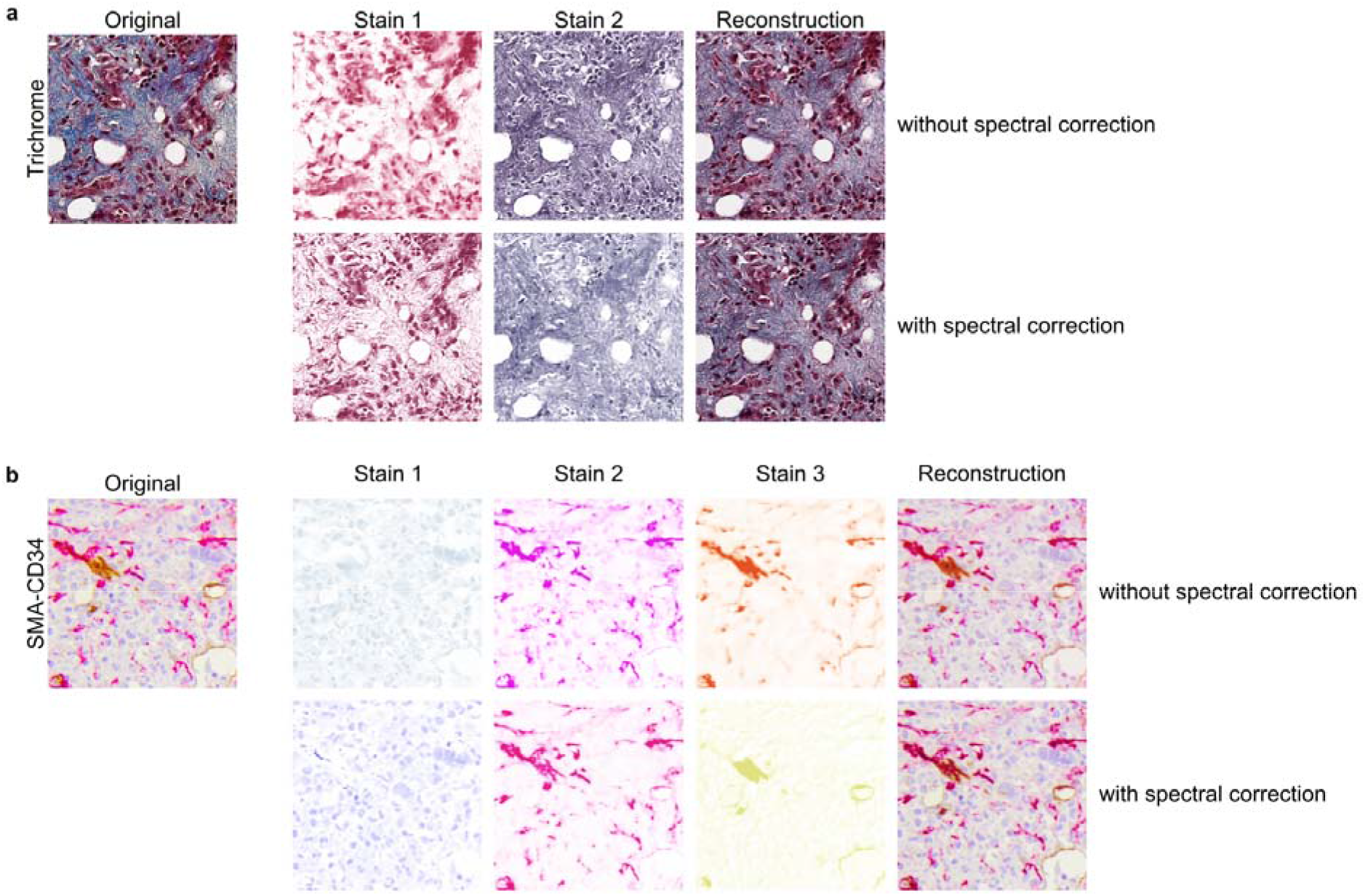
Spectral correction factor improves deconvolution performance. PGDIPS stain deconvolution results of two immunohistochemistry images stained for **a)** Trichrome and **b**) SMA-CD34. The first row of deconvoluted stains and reconstructed images were generated using an abridged PGDIPS model that did not learn the spectral correction factor. The second row shows deconvolution results with spectral correction, which were visually more accurate.

**Extended Data Figure 2:**
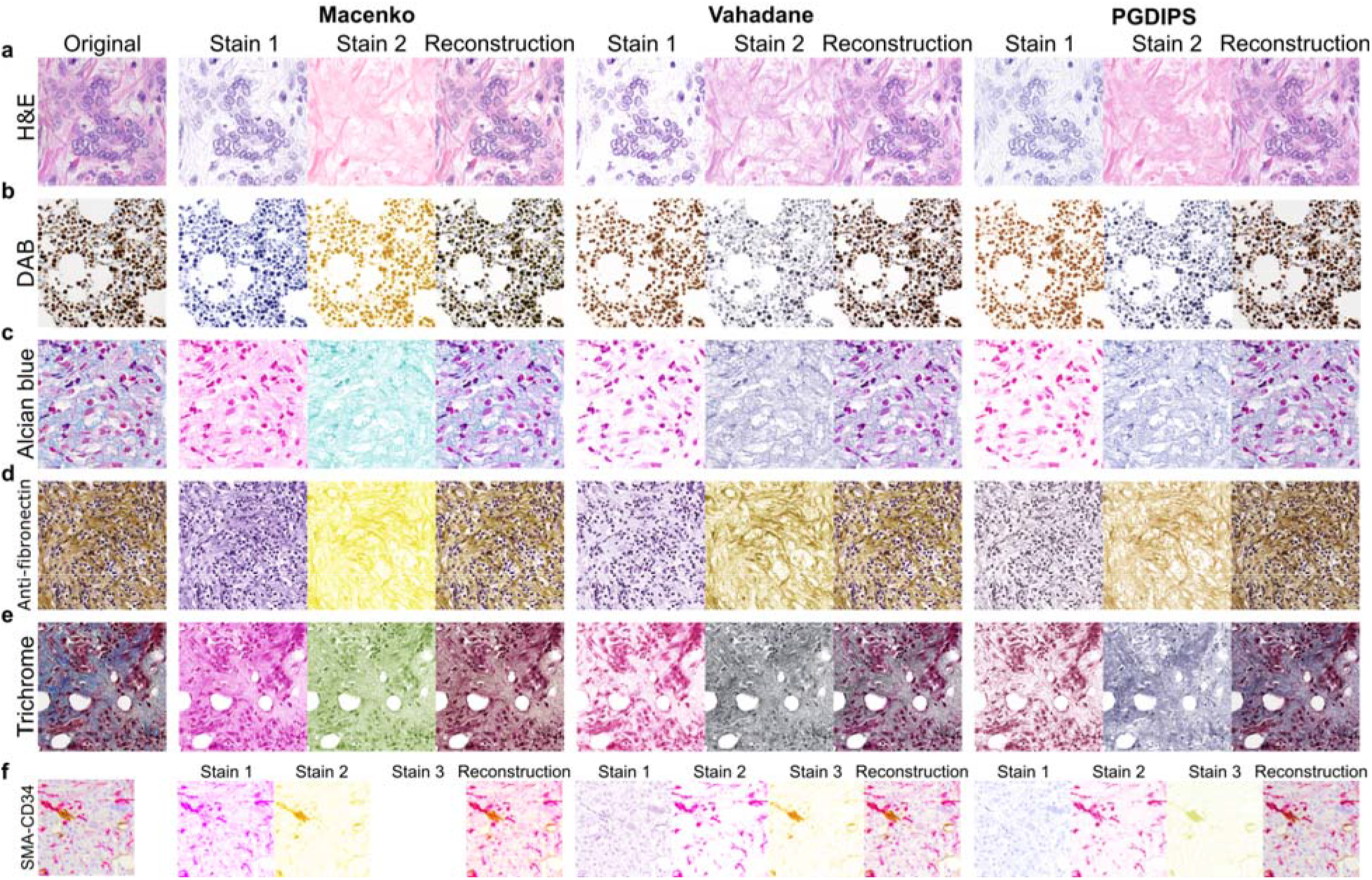
PGDIPS outperforms state-of-the-art algorithms for deconvoluting special stains. Deconvolution results of slides stained with **a**) Hematoxylin and Eosin (H&E) **b**) hematoxylin and 3,3’-diaminobenzidine (DAB) **c**) Alcian Blue **d**) anti-fibronectin antibody **e**) Masson’s Trichrome **f**) SMA-CD34 dual antibodies. Columns are organized to display the target image, the colored concentration maps (matrix multiplication of corresponding stain color vectors and stain concentration maps) and the reconstructed image for Macenko, Vahadane and PGDIPS. PGDIPS achieved visually superior deconvoluted stains with higher fidelity and accuracy while Macenko and Vahadane had poor performance on one or more stain types. Note that it is non-trivial to adapt Macenko *et al*. for three stains so only two stains were deconvoluted by Macenko for SMA-CD34.

**Extended Data Figure 3:**
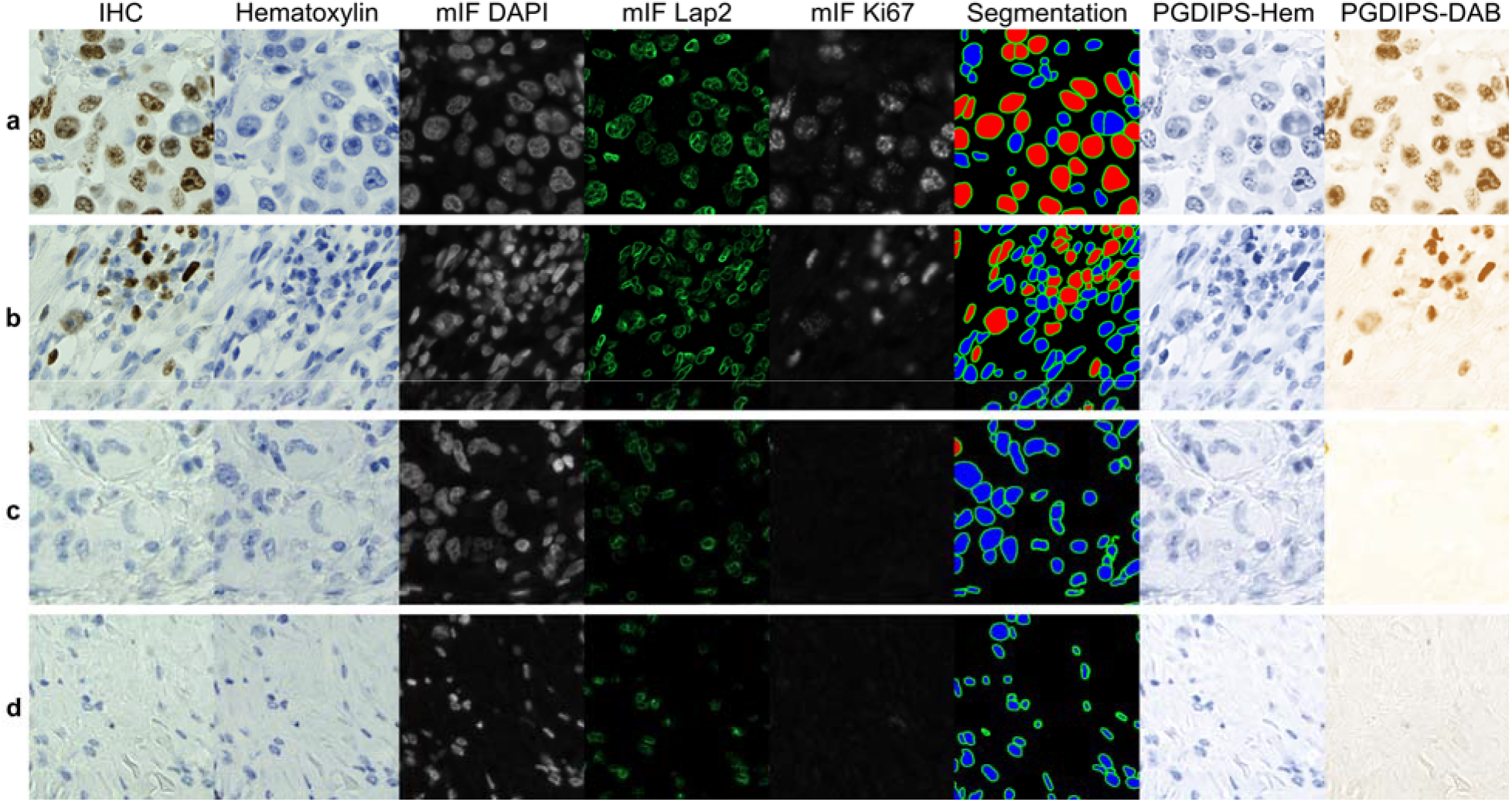
Variations in DAB stain distributions affect deconvolution performance. Example images from the test set of DeepLIIF that have relatively **a**) high **b**) medium **c)** low and **d**) no 3,3’-diaminobenzidine (DAB) staining in immunohistochemistry (IHC) of DAB with hematoxylin as counterstain. Columns from the left show the original DAB image, followed by the co-registered hematoxylin-only stain of the same tissue, immunofluorescence (mIF) of 4’,6-diamidino-2-phenylindole (DAPI), Lap2 and Ki67, cell segmentation provided as part of the DeepLIIF test set (blue cells: DAB-negative, red cells: DAB-positive), and stain deconvolution results by PGDIPS for hematoxylin (Hem) and DAB. Deconvolution was consistent for hematoxylin in **a**), **b**) and **c**) but varied for DAB, especially in **d**) where the model learned a different color for hematoxylin and noise as the second stain instead of DAB (which was absent from the original image). We reason that this explains the larger variations of color vectors estimated in this dataset and the lower long tails of pearson correlations between deconvoluted DAB and mIF Ki67.

**Extended Data Figure 4:**
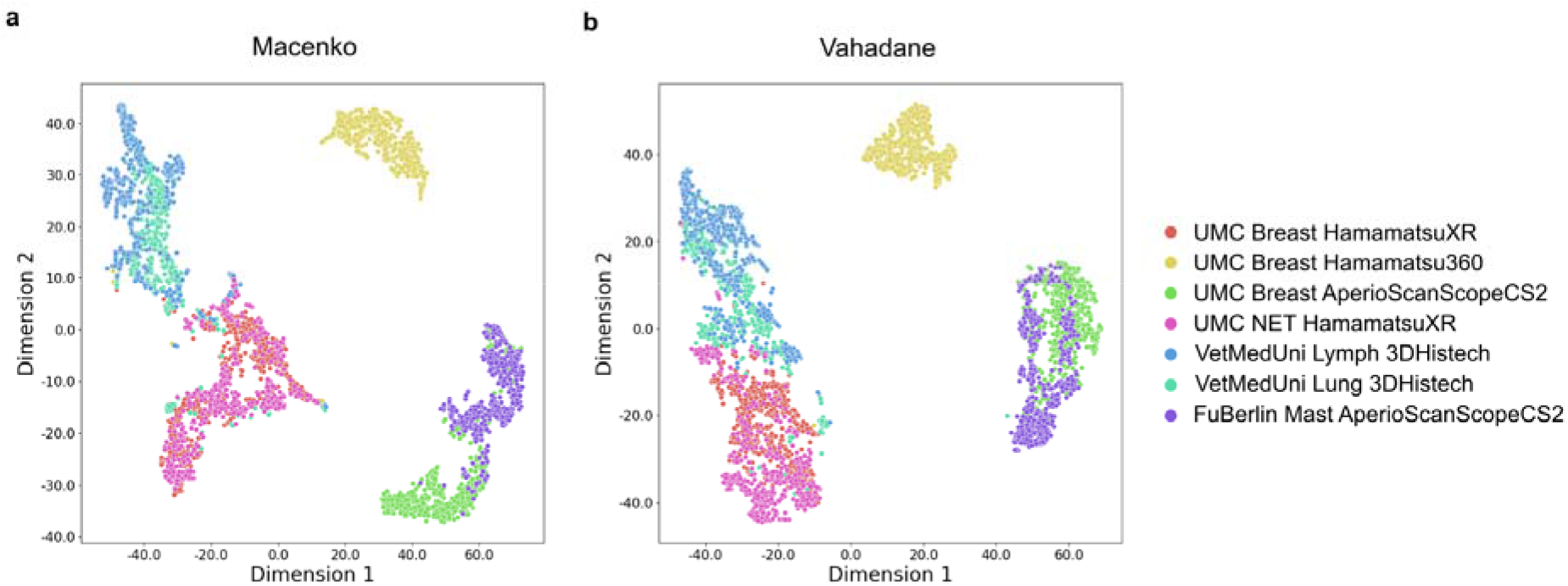
Stain parameter estimation on the MIDOG2022 dataset by Macenko and Vahadane. Estimated stain color vectors of images from MIDOG2022, with dimension reduction and visualization using t-distributed stochastic neighbor embedding (t-SNE), deconvoluted using **a**) Macenko **b**) Vahadane. Both algorithms had difficulty distinguishing images scanned with HamamatsuXR and 3DHistech. n=3186 for both panels. Breast: human breast cancer; NET: human neuroendocrine tumor; Lymph: canine lymphoma; Lung: canine lung cancer; Mast: canine cutaneous mast cell tumor. UMC: University Medical Center Utrecht; VetMedUni: University of Veterinary Medicine Vienna; FuBerlin: Free University of Berlin.

## Supplementary File

**Supplementary Table 1:**
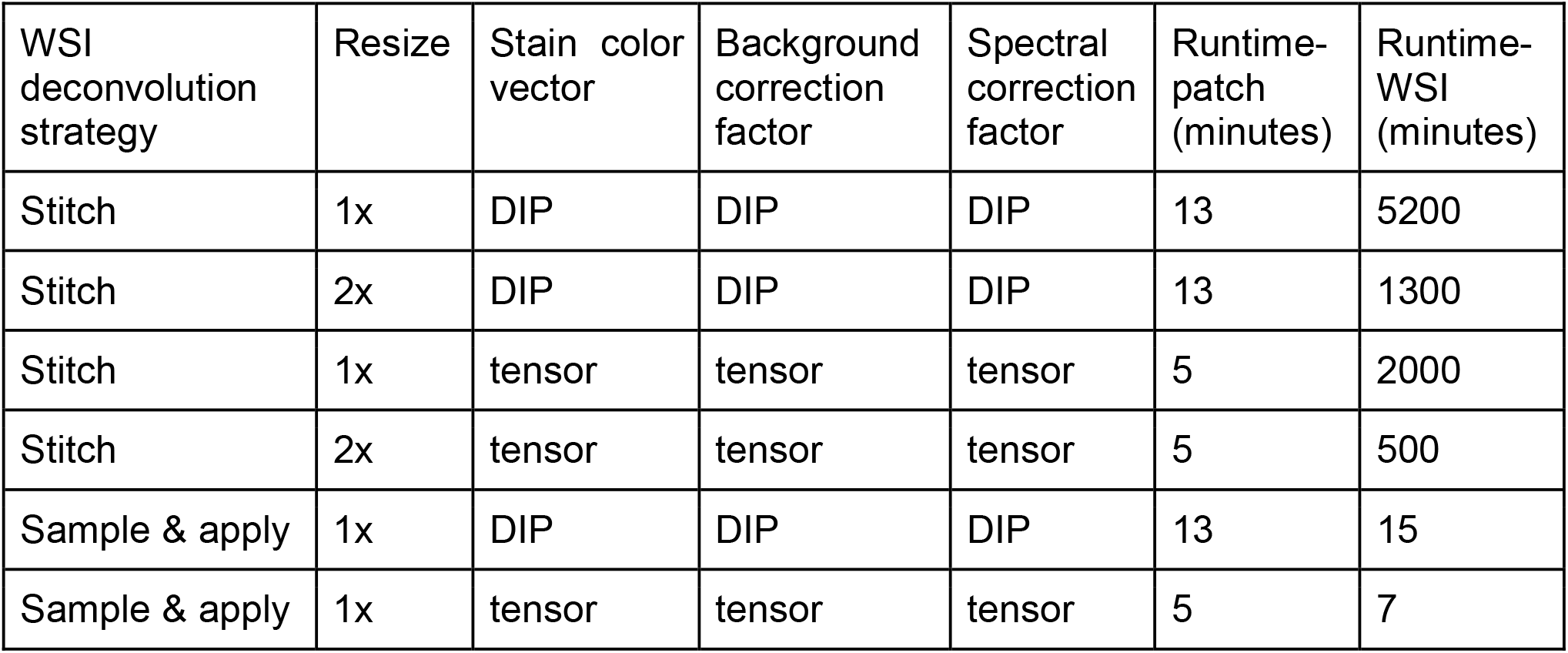
Comparison of whole slide image stain deconvolution strategies.

Strategies for accelerating PGDIPS for the stain deconvolution of a whole slide image (WSI). The baseline strategy is to divide the WSI into smaller patches and perform SD for all patches, and then stitch the deconvolution results into concentration maps the size of the original WSI. Another strategy ‘Sample’ refers to sampling a representative patch from the WSI, using PGDIPS to calculate stain parameters (color vectors, background correction factor and spectral correction factors), and then applying a conventional SD algorithm with the stain parameters to obtain results for the WSI. Resizing the WSI 2x can speed up PGDIPS SD by 4x. In an abridged PGDIPS model where stain parameters are optimized as tensor variables directly instead of using DIPS, the runtime of PGDIPS on a patch can be reduced from 13 minutes to 5 minutes at the cost of a small amount of accuracy. Runtimes were obtained from experiments on an NVIDIA RTX 3080 graphics card and an AMD Ryzen 7 3800X processor, with Vahadane as the conventional SD algorithm, assuming a patch size of 500×500 pixels and a WSI size of 10000×10000 pixels.

